# On the annotation of Paleo- and Mesoproterozoic microfossils as fungi

**DOI:** 10.1101/2024.01.25.577314

**Authors:** Dheeraj Kanaparthi, Marko Lampe, Michel Krings, Bettina Schue, Andreas Klingl, Petra Schwille, Tillmann Lueders

## Abstract

How old is the crown group Fungi? Inferences from phylogenetic and fossil-based studies provided far-apart age estimates ranging between 0.75 to 2.7 billion years old. One important criterion for interpreting Paleo- and Mesoproterozoic microfossils as Fungi is their uncanny morphological resemblance with extant fungi. Here, we demonstrate that bacteria exposed to environmental conditions similar to the paleoenvironmental settings where these presumed fungi lived can spontaneously transform into their protoplasts. These protoplasts exhibit morphologies corresponding to those of presumed fungal microfossils. These observations, together with microfossil chemical composition, pose a serious challenge to interpreting Paleo- and Mesoproterozoic microfossils as fungi. Based on these results, we reiterate that morphology is not a reliable indicator of the phylogeny of microfossils older than 2.5Ga.

---

Molecular clock estimates suggest that fungi are an ancient lineage of organisms that must have evolved relatively early in geological time, probably in the Mesoproterozoic (1.6-1.0 Ga)^1^. The earliest fungi were thought to have been phagotrophic marine organisms that lacked a chitinous cell wall during their trophic phase ^2,3^. Early diverging fungal lineages that still exist today, like the Aphelida, Microsporidia, and Cryptomycota (Genus *Rozella*), still live a similar life cycle and thrive as parasites ^4^. Geologically younger species of fungi were thought to have evolved either by reductive genome evolution (endoparasite fungi) ^5,6^ or by developing a saprophytic lifestyle dependent on plant-derived organic carbon (ectosymbiotic fungi) ^7,8^. This interdependency between plants and fungi was thought to have led to the co-evolution of fungi together with plants. Such symbiotic associations were hypothesized to have given rise to members of the sub-kingdom Dikarya ^9^.

The sequence of events described above was postulated from studying the phylogeny and genomes of the extant fungi and, to a limited extent, corroborated by the fungal fossil record^10^. The oldest bonified fossils of Fungi are reported from the Ordovician ^11^. The oldest preserved fungal and plant microfossils together were reported from Rainy Chert ^12–14^. Given the close interaction observed between fungal and plant microfossils in Rainy chert, they were thought to have been in a symbiotic relationship with early land plants. Such symbiotic associations were hypothesized to have facilitated the colonization of land by plants ^15,16^.

The past few years have seen the discovery of several microfossils attributed to fungi that were not in accordance with the generally accepted evolutionary timeline. These studies propose that higher fungi belonging to Dikarya could have evolved as early as 2.4 billion years ago ^17^ and colonized terrestrial environments at least by 1 Ga ^18,19^. ∼1.2 billion and ∼500 million years earlier than the currently accepted time period ^10,11^. Resolving this discrepancy is important for our understanding of fungal evolution. Moreover, understanding the timeline of fungi is inextricably linked to understanding the evolutionary timeline of other eukaryote lineages ^1,20^, the evolution of land plants ^9^, and the geological implications of land colonization by plants ^21,22^. Here, we attempt to resolve the above-mentioned discrepancies regarding the origin of Fungi.

One key factor that has fostered the attribution of *Paleo- and Mesoproterozoic* microfossils from Ongeluk formation (_∼_ 2.4 Ga) ^17^, Gunflint iron formations (_∼_ 2Ga) ^23^, Grassy Bay formation (_∼_ 1Ga) ^18^, and Lakhanda formation (_∼_ 1Ga) ^19^, is their similarity in size and basic morphology to extant fungi. However, we have previously shown that bacteria could transform into their protoplast state under some environmental conditions ^24–26^, and cell wall could have evolved no earlier than ∼2 Ga (this manuscript is under preparation). Based on these results, we argue that morphology is not a reliable indicator in the identification of fungal microfossils. Proterozoic microfossils attributed to Fungi could as well be bacterial protoplasts. Below, we present our argument from a geological, morphological, chemical, and biological perspective.

One of the environmental factors that contributes to the loss of bacterial cell wall is the concentration and nature of salt in the environment. Several phylogenetic groups of bacteria have been shown to spontaneously transform into their protoplast state when exposed to salinities above 5% w/v (R). Such salinity levels were prevalent in the paleo-environments in which the organisms interpreted as fungi inhabited. Geologically, all the source strata mentioned above were deposited in shallow marine paleo-environments ^27^. Evidence for periodic evaporation, leading to elevated salt concentrations, was reported from these sites. As an indication of this process, several sites within the Shaler formation were known to have formed desiccation cracks and evaporation mounds ^28^. The salinities of these sites were known to fluctuate between 2-4 times the current ocean salinities. Similarly, other sites, Ongeluk formation (part of Barberton Green Belt) ^27^, Lakhanda formation ^29^, and Gunflint iron formations ^30^, were also known to have experienced higher salinities.

Given the higher salinities of these sites, we cultured Gram-negative *Rhodobacter sphaeroides* in nutrient broth (NB) amended either with 6-12% DSS or NaCl (wt/vol). Cultivation bottles were incubated at 30^0^C under static conditions. Cells in these incubations were observed under a microscope at regular intervals. Over the course of these incubations, *R.sphaeroides* grew as a biofilm attached to the bottom of the cultivation flask and transformed into its protoplast state (*RS-P*), even in the absence of cell wall degrading enzymes like lysozyme or antibiotics like Penicillin G. This transition was evident from the morphological similarities with cells from our previous studies (Figure 1). Morphologies of such cells were remarkably similar to that of microfossils reported from the Ongeluk, Gunflint, Lakhanda formation, and Grassy Bay formations.

**Fig 1.**
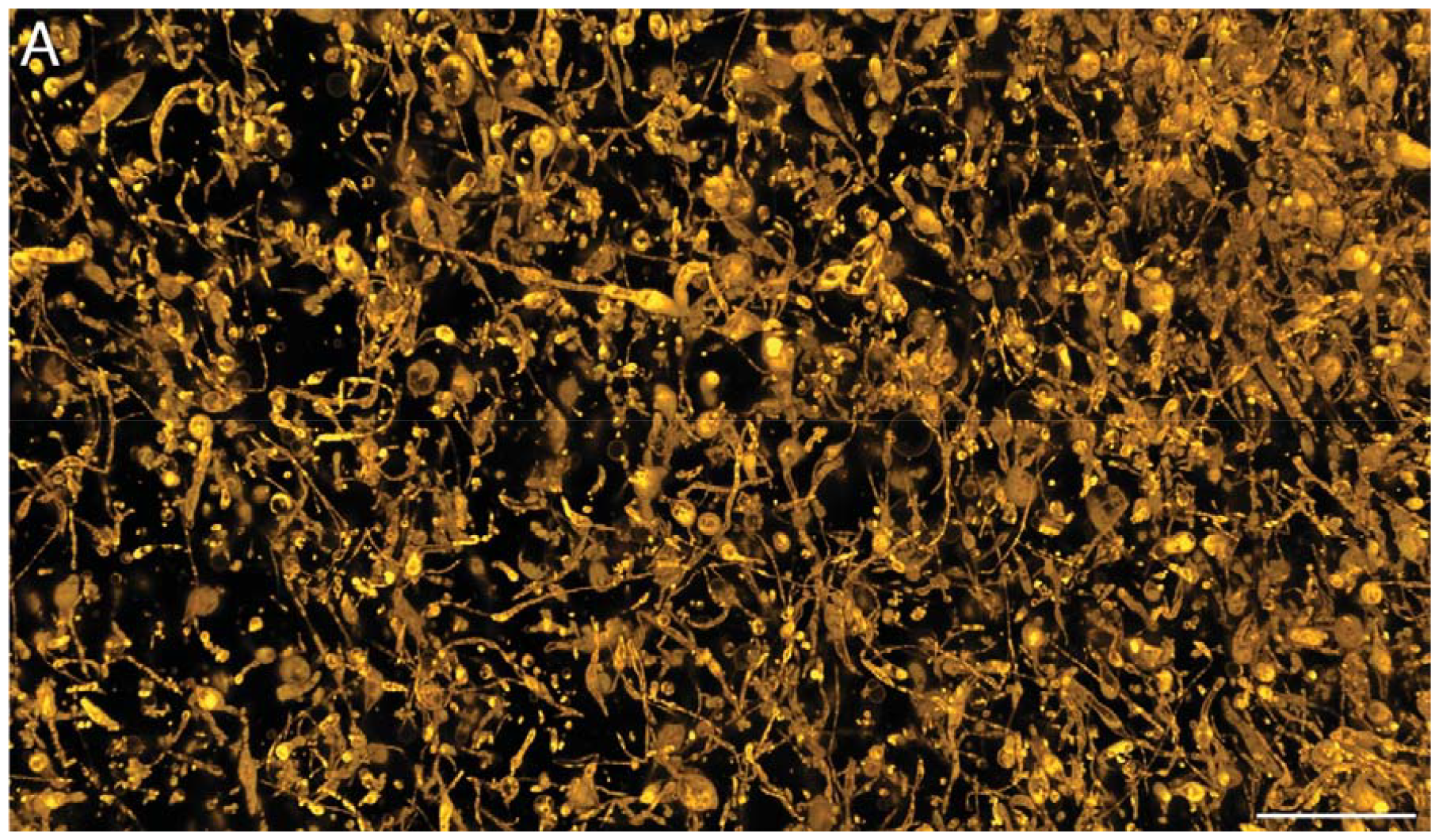
Protoplasts of R. sphaeroides. Image A is the confocal microscope image of *RS-P* (Scale bar: 20 µm). Cells in these images were stained with universal membrane stain, FM™5-95 (yellow). Thin or thick tubular cells with bulbous structures can be seen among *RS-P* cells.

Microfossils reported from the Ongeluk Formation occur in the form of filamentous cells encrusted in calcite ^17^. Chronologically, these are the oldest microfossils to date that have been interpreted as fungi. Justifiably, morphological features exhibited by these microfossils are distinctively fungal. Nevertheless, *RS-P* exhibited remarkably similar morphologies (Fig. 1&2). Like the microfossils, *RS-P* grows as a biofilm attached to surfaces and forms an entangled web of filamentous cells (Fig. 1&2). Filamentous *RS-P* cells formed loops of different diameters that resemble the pattern described in the Ongeluk Formation (Fig. S1-S3). We also observed Y and T-type branching of the filamentous *RS-P* cells resemble the branching reported from Ongeluk fossils (Fig. 2, S4 & S5). Our observations from *RS-P* were in accordance with the proposition that Y-type branching could have been the true branching compared to T-type branching, which could have been formed by the juxtaposition of two filamentous cells. In support of this claim, we observed cytoplasmic continuity within the Y-junctions of *RS-P* (Fig. 3 & S4). Local swelling or inflation 5-10 µm and occurring at both terminal and intercalary positions were observed in the bacterial filaments. They closely resemble the structures reported from the microfossils (Fig. 1&2). Based on the similarities in morphology and surface texture, we presume that the basal film-like structure observed within these microfossils could have been the membrane debris observed in *RS-P* (Fig. S6 & S7).Similar structures with entangled filamentous cells were often in *RS-P*’s incubations.

**Fig 2.**
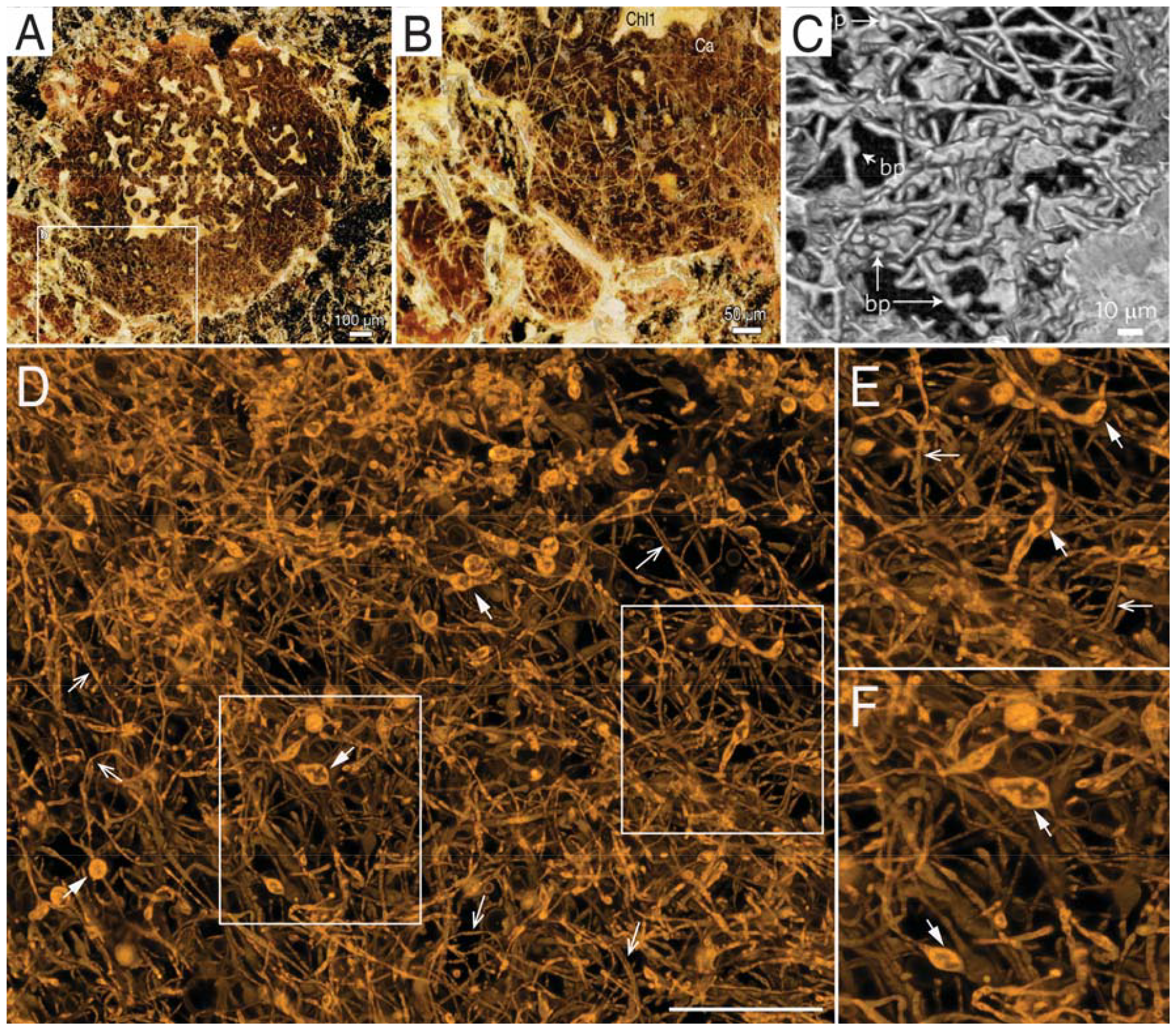
Morphological comparison of Ongeluk formation microfossils with *RS-P*. Images A-C are microfossils reported from Ongeluk formation. Image D is a confocal microscope image of *RS-P* (Scale bar: 50 µm) (also see Fig. S1). Cells in these images were stained with universal membrane stain, FM™5-95 (yellow). Arrows in image C point to bulbous protrusions within the filaments. Closed arrows in image D point to similar protrusions in *RS-P*. Open arrows in image D point to loops formed by *RS-P* (also see Fig. S2 & S3). Images E & F are the magnified regions of the D.

**Fig 3.**
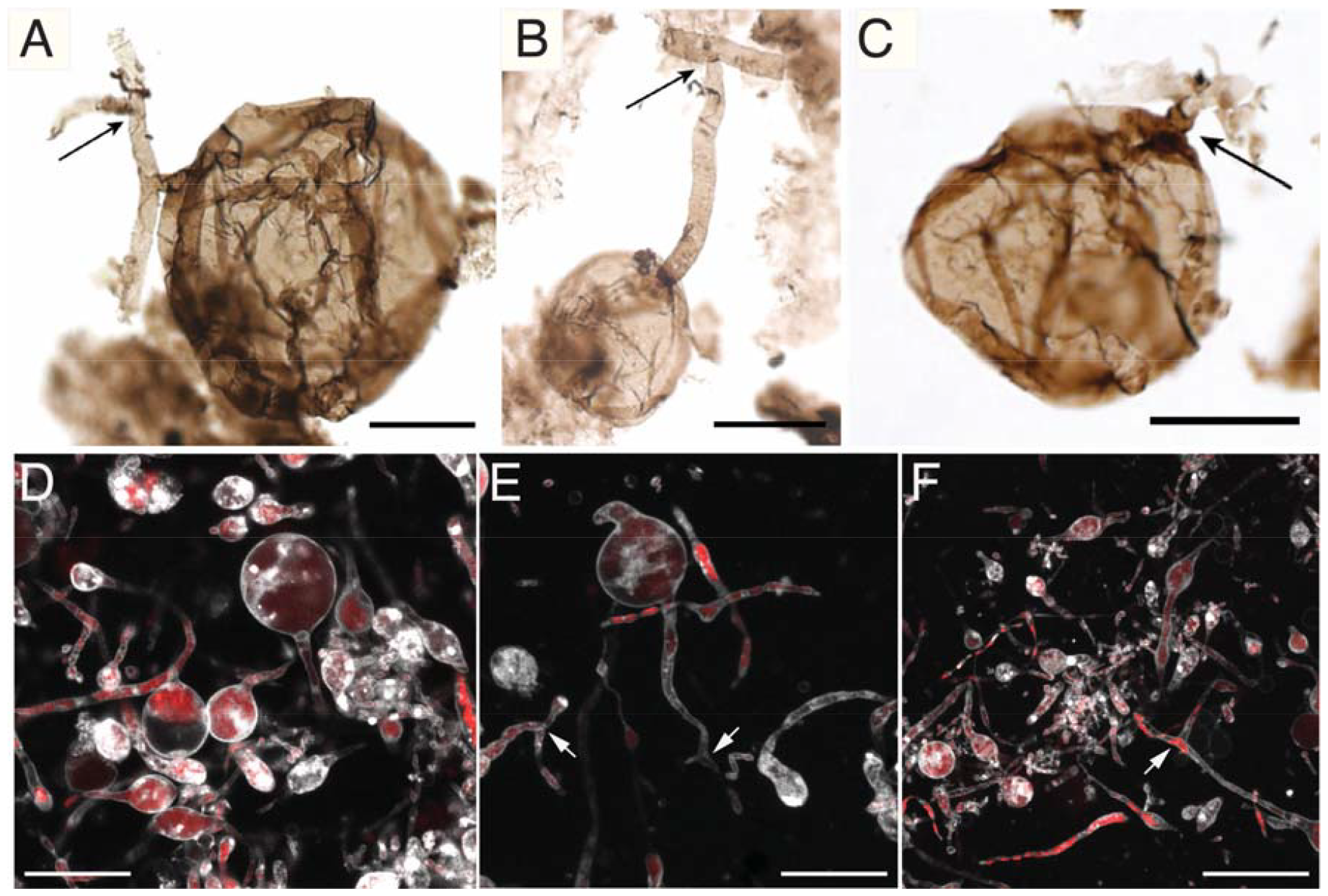
Morphological comparison of Grassy Bay formation microfossils with *RS-P*. Images A-C are microfossils reported from the Grassy Bay formation. Image D is a confocal microscope image of *RS-P* (Scale bar: 10 µm) (also see Fig. S14-16). Cells in these images were stained with universal membrane stain, FM™5-95 (white), and DNA stain, PicoGreen™ (red). Arrows in images A-C point to branching within filamentous cells. Arrows in images D&F point to a similar branching pattern in *RS-P* (also see Fig. S4).

Microfossils reported from Gunflint iron formations, Lakhanda Formation, and Grassy Bay Formation are also morphologically similar to *RS-P*. All morphological features of these fossils that were assumed to be distinctively fungal, including spherical bulbous structures with lateral or vertical septa resembling conidia (Fig. S8-S10), intertwined filamentous cells resembling mycelia (Fig. S11), and the terminal intercalary inflations similar to chlamydospores, have also been observed in *RS-P* (Fig. 3). Different developmental stage od “germinating” conidia-like structures in *RS-P* are shown in Fig. S13. Morphological comparison of Grassy Bay formation microfossils with *RS-P* is presented in Fig. S14 & S16.

Along with their morphological resemblance, earlier studies claimed the presence of chitin within Grassy Bay microfossil biomass. Detection of chitin by FTIR analysis was argued as an indication that these microfossils were Fungi. We disagree with this interpretation. Identification of chitin relied on the observation of absorption bands between 900 and 1300cm^-1^. While the spectra of chitin and chitosan are dominated by complex multiplets between 900 and 1300 cm^-1^, which arise from the absorption of C-O bonds in the carbohydrates, they are not specific for either compound. Most organic and biological compounds show absorption in this spectral region (Fig. 4) ^31^. Compounds similar to chitin and chitosan, such as peptidoglycan, preparations of *Cantharellus cibarius*, and even the bacterial protoplasts devoid of a peptidoglycan cell wall showed a similar band pattern (Fig. 4), implying that these absorption bands could be due to carbohydrates within the cell or associated with the cell membrane, rather than a specific marker for chitin. One other possibility for the observed adsorption pattern could also be contamination from silica, silicates, and phosphate-containing minerals that were closely associated with the microfossil, which are known to exhibit multiplets between 900 and 1300 cm^-1 32^. Hence, we argue that band patterns in this spectral region are not a reliable indication of chitin, especially if the samples represent multicomponent mixtures.

**Figure 4.**
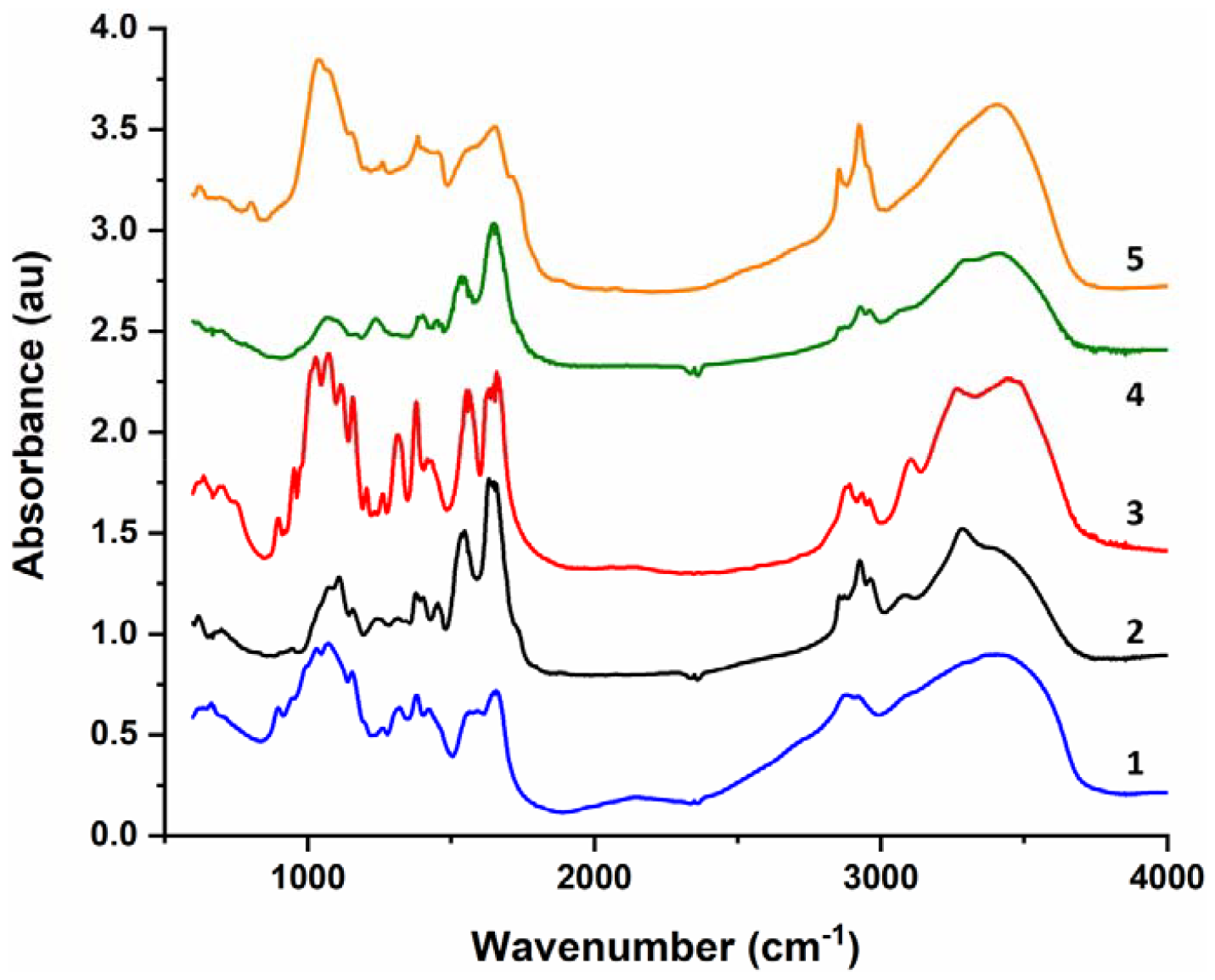
Comparison of FTIR spectra of reference biopolymer and selected organisms. 1: chitosan; 2: peptidoglycan; 3: chitin; 4: *RS-L*; 5: Cantharellus cibarius.

Absorption of chitin between 2800 and 3300 cm^-1^ shows a characteristic sequence of bands (1880, 2935, 2960, 3100, 3260 cm^-1^). Spectra from Grassy Bay microfossils attributed to vestigial chitin instead show two strong bands at 2850 and 2920 cm^-1 18^. These bands are characteristic of long alkyl chains (triglycerides, fatty acids, hydrocarbon fractions, phospholipids), rather than polysaccharides or proteins ^31^. This is in accordance with our proposition that these microfossils were, most likely, bacteria in their protoplast state and that the microfossil biomass, most likely, was derived from their lipid membrane. The wrinkly nature of the microfossils also supports our assumption that these structures were derived from a flexible membrane rather than a rigid cell wall ^33^.

Biosynthetic pathways of both peptidoglycan in bacteria, chitin, and cell wall glycoproteins in eukaryotic cells are remarkably similar ^34,35^. Both pathways involve the synthesis of UDP-N-acetylglucosamine, subsequent addition of amino acids to UDP-N-acetylglucosamine, translocation across the cytoplasmic membrane using a lipid carrier, and polymerization on the cell surface. Several enzymes involved in this process, like the *marY* gene in *E.coli*, the ALG7 gene in yeast, and the *GTP* gene in hamsters, have been shown to have evolved from a common ancestor and share considerable sequence similarity ^36,37^. Given the highly conserved nature of these pathways, it has been hypothesized that these pathways first evolved in prokaryotes for peptidoglycan synthesis and subsequently underwent diversification to facilitate a plethora of glycosylation reactions in eukaryotes ^38,39^. The earliest known evidence of a bacterial cell wall comes from the Gunflint Iron Formations (approx. 2Ga). This suggests that eukaryotes with a carbohydrate-containing cell wall, like fungi, could only have evolved much later. This is not in accordance with the interpretation of the Ongeluk Formation microfossils as fungi.

Evolutionary relationships among different organisms can be assessed by constructing phylogenetic trees. Although genome-based analysis provides information regarding the shared ancestor and evolutionary variation between different lineages, translating phylogenetic variation to geological time requires corroboration from the fossil record. The evolutionary history of fungi remains highly debated among scholars as both results from phylogenetic analyses and the interpretations of the fossil record were not in tune with each other. Earlier studies demonstrated that one possible reason for these discrepancies could be due to the misinterpretation of some microfossils as fungi ^10^. The results we presented above corroborate this proposition. Together with the results from our earlier work, we propose that morphology is not a reliable indicator for predicting the taxonomy of microfossils, especially the microfossils older than 2Ga. Based on our results, we also argue for the imposition of a hardbound constraint on the evolution of bacterial cell wall at 2 Ga. By extension, all eukaryotes with a carbohydrate-based cell wall could only have evolved much later.

## References

1. Parfrey, L. W., Lahr, D. J. G., Knoll, A. H. & Katz, L. A. Estimating the timing of early eukaryotic diversification with multigene molecular clocks. Proc. Natl. Acad. Sci. U. S. A. (2011) doi:10.1073/pnas.1110633108.

2. Del Campo, J. & Ruiz-Trillo, I. Environmental survey meta-analysis reveals hidden diversity among unicellular opisthokonts. Mol. Biol. Evol. (2013) doi:10.1093/molbev/mst006.

3. Cavalier-Smith, T. Megaphylogeny, cell body plans, adaptive zones: Causes and timing of eukaryote basal radiations. in Journal of Eukaryotic Microbiology (2009). doi:10.1111/j.1550-7408.2008.00373.x.

4. James, T. Y. et al. A molecular phylogeny of the flagellated fungi (Chytridiomycota) and description of a new phylum (Blastocladiomycota). Mycologia (2006) doi:10.3852/mycologia.98.6.860.

5. James, T. Y. et al. Shared signatures of parasitism and phylogenomics unite cryptomycota and microsporidia. Curr. Biol. (2013) doi:10.1016/j.cub.2013.06.057.

6. Haag, K. L. et al. Evolution of a morphological novelty occurred before genome compaction in a lineage of extreme parasites. Proc. Natl. Acad. Sci. U. S. A. (2014) doi:10.1073/pnas.1410442111.

7. Delaux, P. M. et al. Algal ancestor of land plants was preadapted for symbiosis. Proc. Natl. Acad. Sci. U. S. A. (2015) doi:10.1073/pnas.1515426112.

8. Chang, Y. et al. Phylogenomic analyses indicate that early fungi evolved digesting cell walls of algal ancestors of land plants. Genome Biol. Evol. (2015) doi:10.1093/gbe/evv090.

9. Feijen, F. A. A., Vos, R. A., Nuytinck, J. & Merckx, V. S. F. T. Evolutionary dynamics of mycorrhizal symbiosis in land plant diversification. Sci. Rep. (2018) doi:10.1038/s41598-018-28920-x.

10. Berbee, M. L., James, T. Y. & Strullu-Derrien, C. Early Diverging Fungi: Diversity and Impact at the Dawn of Terrestrial Life. Annu. Rev. Microbiol. (2017) doi:10.1146/annurev-micro-030117-020324.

11. Redecker, D., Kodner, R. & Graham, L. E. Glomalean fungi from the Ordovician. Science (2000) doi:10.1126/science.289.5486.1920.

12. Krings, M., Taylor, T. N. & Dotzler, N. Fossil evidence of the zygomycetous fungi. Persoonia: Molecular Phylogeny and Evolution of Fungi Preprint at 10.3767/003158513X664819 (2013).

13. Krings, M., Taylor, T. N. & Dotzler, N. Fossil evidence of the zygomycetous fungi. Persoonia: Molecular Phylogeny and Evolution of Fungi vol. 30 1–10 Preprint at 10.3767/003158513X664819 (2013).

14. Taylor, T. N., Remy, W. & Hass, H. FUNGI FROM THE LOWER DEVONIAN RHYNIE CHERT: CHYTRIDIOMYCETES. Am. J. Bot. (1992) doi:10.1002/j.1537-2197.1992.tb13726.x.

15. Simon, L., Bousquet, J., Lévesque, R. C. & Lalonde, M. Origin and diversification of endomycorrhizal fungi and coincidence with vascular land plants. Nature (1993) doi:10.1038/363067a0.

16. Humphreys, C. P. et al. Mutualistic mycorrhiza-like symbiosis in the most ancient group of land plants. Nat. Commun. (2010) doi:10.1038/ncomms1105.

17. Bengtson, S. et al. Fungus-like mycelial fossils in 2.4-billion-year-old vesicular basalt. Nat. Ecol. Evol. (2017) doi:10.1038/s41559-017-0141.

18. Loron, C. C. et al. Early fungi from the Proterozoic era in Arctic Canada. Nature (2019) doi:10.1038/s41586-019-1217-0.

19. German, T. N. & Podkovyrov, V. N. The role of cyanobacteria in the assemblage of the Lakhanda Microbiota. Paleontol. J. (2011) doi:10.1134/S0031030111020079.

20. Douzery, E. J. P., Snell, E. A., Bapteste, E., Delsuc, F. & Philippe, H. The timing of eukaryotic evolution: Does a relaxed molecular clock reconcile proteins and fossils? Proc. Natl. Acad. Sci. U. S. A. (2004) doi:10.1073/pnas.0403984101.

21. Lenton, T. M., Crouch, M., Johnson, M., Pires, N. & Dolan, L. First plants cooled the Ordovician. Nat. Geosci. (2012) doi:10.1038/ngeo1390.

22. Lenton, T. M. et al. Earliest land plants created modern levels of atmospheric oxygen. Proc. Natl. Acad. Sci. U. S. A. (2016) doi:10.1073/pnas.1604787113.

23. McMenamin, M. A. S., Curtis-Hill, A., Rabinow, S., Martin, K. & Treloar, D. A ‘Giant Microfossil’ from the Gunflint Chert and its Implications for Fungal and Eukaryote Origins. (2019) doi:10.20944/PREPRINTS201909.0287.V1.

24. Ramijan, K. et al. Stress-induced formation of cell wall-deficient cells in filamentous actinomycetes. Nat. Commun. 9, (2018).

25. Kanaparthi, D. et al. On the reproductive mechanism of Gram-negative protocells. bioRxiv 2021.11.25.470037 (2021) doi:10.1101/2021.11.25.470037.

26. Kanaparthi, D. et al. On the reproductive mechanisms of Gram-positive protocells. bioRxiv 2021.11.25.470039 (2021) doi:10.1101/2021.11.25.470039.

27. De Ronde, C. E. J., Channer, D. M. D., Faure, K., Bray, C. J. & Spooner, E. T. C. Fluid chemistry of Archean seafloor hydrothermal vents: Implications for the composition of circa 3.2 Ga seawater. Geochim. Cosmochim. Acta (1997) doi:10.1016/S0016-7037(97)00205-6.

28. Mathieu, J.-P. Characteristics of diagenetic fluids affecting two major carbonate units on Victoria Island, Northwest Territories. (2014).

29. Podkovyrov, V. N. Mesoproterozoic Lakhanda Lagerstätte, Siberia: Paleoecology and taphonomy of the microbiota. Precambrian Res. 173, 146–153 (2009).

30. Licari, G. R. & Cloud, P. E. Reproductive structures and taxonomic affinities of some nannofossils from the Gunflint Iron Formation. Proc. Natl. Acad. Sci. 59, 1053–1060 (1968).

31. Naumann, D. FT-infrared and FT-Raman spectroscopy in biomedical research. Appl. Spectrosc. Rev. 36, (2001).

32. Goodman, A. et al. Investigating the role of water on CO2-Utica Shale interactions for carbon storage and shale gas extraction activities – Evidence for pore scale alterations. Fuel 242, (2019).

33. Latgé, J. P. The cell wall: A carbohydrate armour for the fungal cell. Mol. Microbiol. (2007) doi:10.1111/j.1365-2958.2007.05872.x.

34. Bugg, T. From peptidoglycan to glycoproteins: Common features of lipid-linked oligosaccharide biosynthesis. FEMS Microbiol. Lett. (1994) doi:10.1016/0378-1097(94)90425-1.

35. Helenius, A. & Aebi, M. Roles of N-linked glycans in the endoplasmic reticulum. Annu. Rev. Biochem. (2004) doi:10.1146/annurev.biochem.73.011303.073752.

36. Ikeda, M., Wachi, M., Jung, H. K., Ishino, F. & Matsuhashi, M. The Escherichia coli mraY gene encoding UDP-N-acetylmuramoyl-pentapeptide:undecaprenyl-phosphate phospho-N-acetylmuramoyl-pentapeptide transferase. J. Bacteriol. (1991) doi:10.1128/jb.173.3.1021-1026.1991.

37. Zhu, X. & Lehrman, M. A. Cloning, sequence, and expression of a cDNA encoding hamster UDP-GlcNAc:dolichol phosphate N-acetylglucosamine-1-phosphate transferase. J. Biol. Chem. (1990) doi:10.1016/s0021-9258(18)77293-1.

38. Samuelson, J. et al. The diversity of dolichol-linked precursors to Asn-linked glycans likely results from secondary loss of sets glycosyltranferases. Proc. Natl. Acad. Sci. U. S. A. (2005) doi:10.1073/pnas.0409460102.

39. Wacker, M. et al. N-linked glycosylation in Campylobacter jejuni and its functional transfer into E. coli. Science (2002) doi:10.1126/science.298.5599.1790.

