## Supplementary materials for "On the annotation of Paleo- and Mesoproterozoic microfossils as fungi"

Materials and Methods

*Rhodobacter sphaeroides* (DSM159) was obtained from DMSZ (Braunschweig, Germany).

Cells were initially cultured in nutrient broth (NB) and subsequently transferred to half-strength nutrient broth (NB) containing 8% Dead Sea salt (8%DSS-NB).

Transformation to their protoplast state (*RS-P*) was confirmed by a change in cell morphology. Over the next 3-4 weeks, the cells were repeatedly sub-cultured into fresh media to test the stability of these protoplasts. For subsequent experiments, protoplasts are grown in their respective media with DSS concentrations ranging between 5-10%.

Morphology *RS-P* under different growth conditions was determined using a Carl Zeiss Axioskop 2plus microscope equipped with Axiocam HRm (Carl Zeiss, Germany) or Leica DSF9000 (Leica, Germany) cameras. TEM was carried out using a Zeiss EM 912 with an integrated OMEGA filter, as described before.

STED microscopy was performed with an inverted TCS SP8 STED 3X microscope (Leica Microsystems, Mannheim Germany) using an 86x/1.2 NA water immersion objective (Leica HC PL APO CS2 - STED White). Fluorophores were excited with either 488, 561nm, 594nm, or 633m laser light derived from an 80 MHz pulsed White Light Laser (Leica Microsystems, Mannheim Germany). For stimulated emission, either a pulsed 775 nm laser or a 592nm CW laser (Leica Microsystems, Mannheim, Germany) was used depending on the fluorophore. The emitted fluorescence light was collected with Hybrid Detectors (HyD, Leica Microsystems, Mannheim Germany) using appropriate emission band pass-filter setting and typically a gate of 0.3ns-6.0ns for depletion with the 775nm laser or 1.0-6.0ns with the 592nm laser. Images were recorded in photon counting mode and line accumulation. Image deconvolution was performed on selected images and movies with Huygens Professional (version 16.10.1p2, Scientific Volume Imaging, Hilversum, The Netherlands).

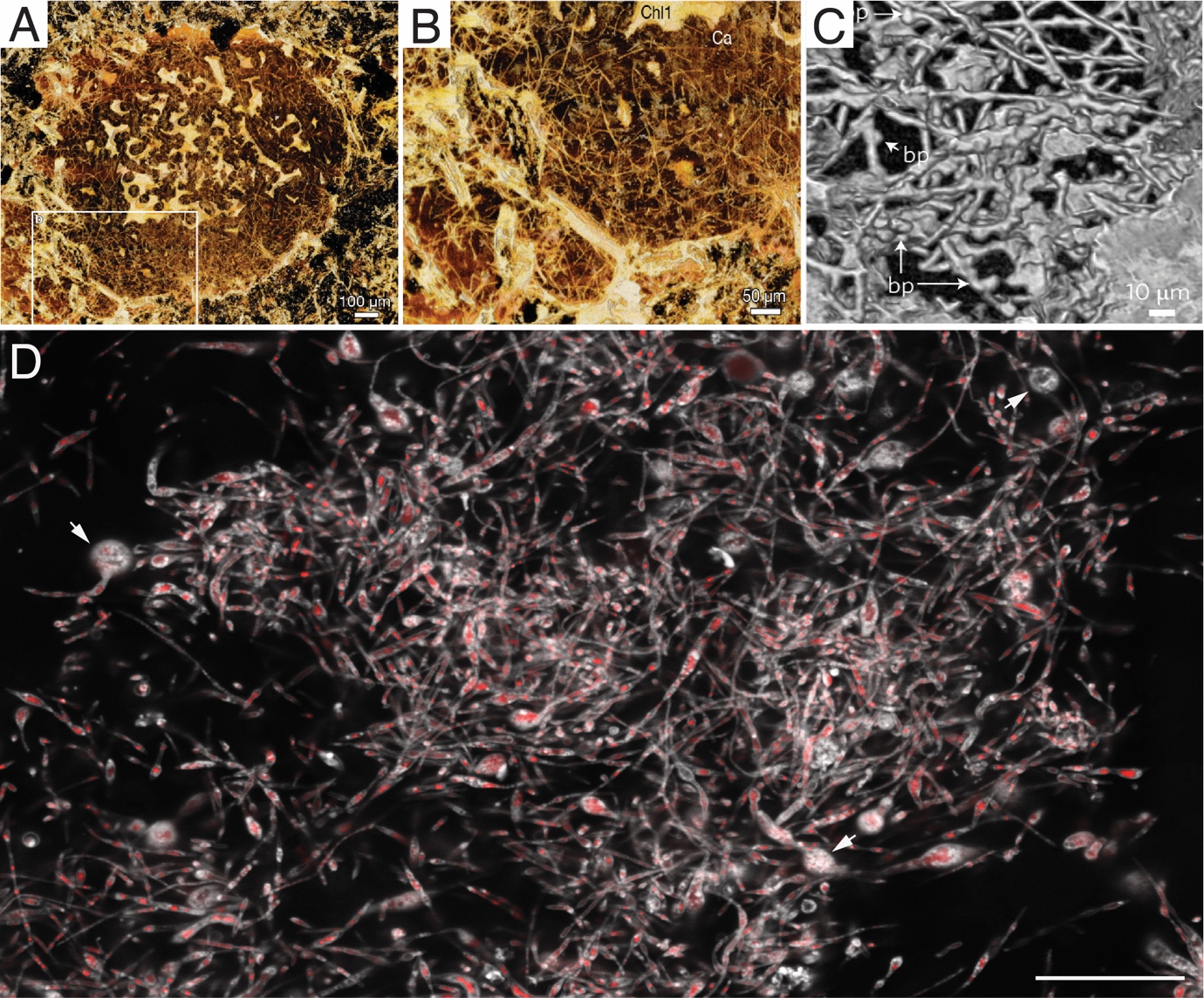

**Fig S1. Morphological comparison of Ongeluk formation microfossils with *RS-P*.**

Images A-C are microfossils reported from Ongeluk formation. Image D is a confocal microscope image of *RS-P* (Scale bar: 20 µm). Cells in these images were stained with universal membrane stain, FM^TM^5-95 (white), and DNA stain, PicoGreen^TM^ (red).

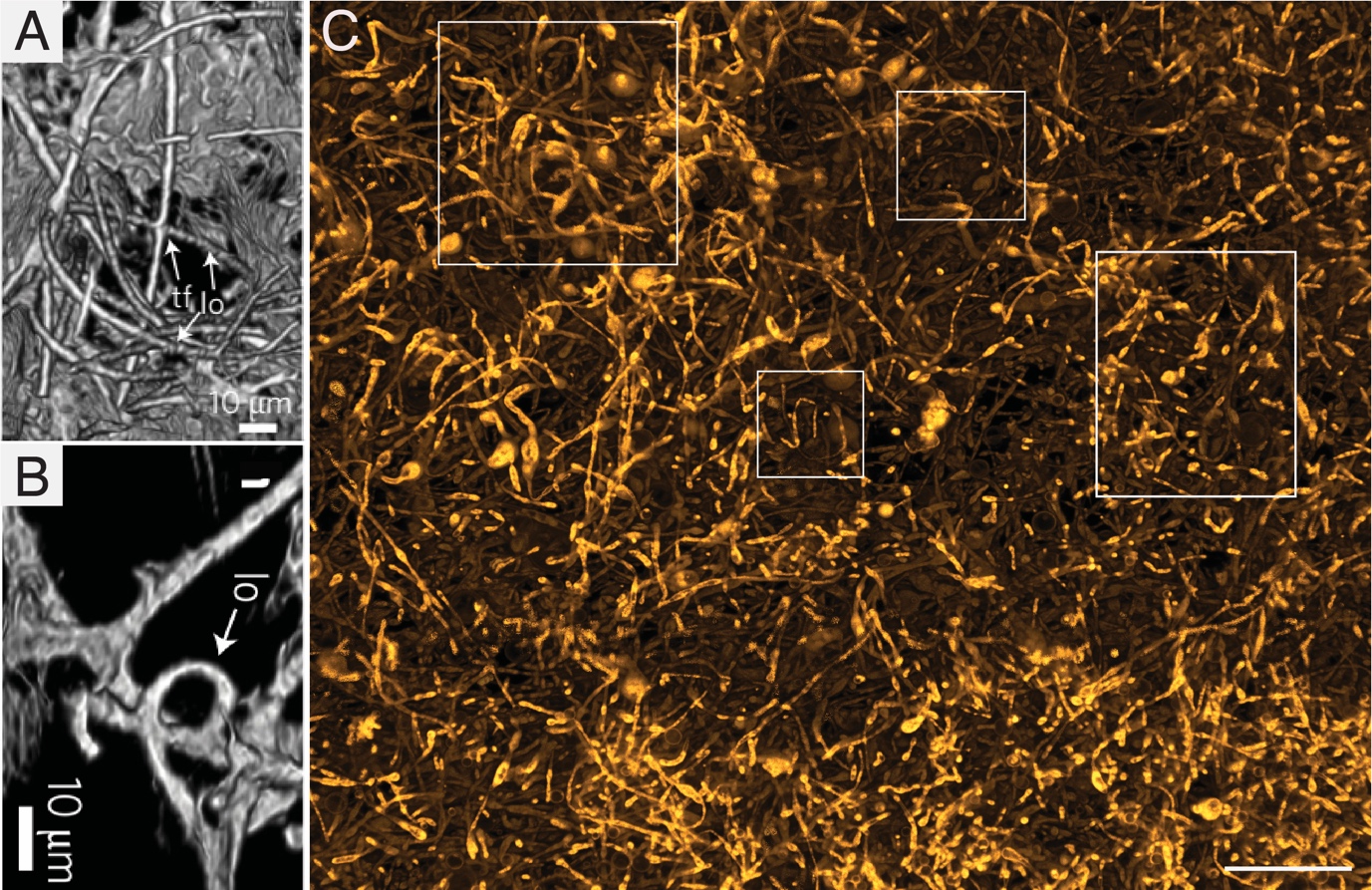

**Fig S2. Morphological comparison of Ongeluk formation microfossils with *RS-P*.**

Images A & B are microfossils reported from Ongeluk formation. Image C is a confocal microscope image of *RS-P* (Scale bar: 20 µm). Cells in these images were stained with universal membrane stain, FM^TM^5-95 (yellow). Arrows (lo) in images A & B point to loops formed by the filamentous microfossils. Boxed regions within image C contain filaments exhibiting a similar loop formation (see Fig S3).

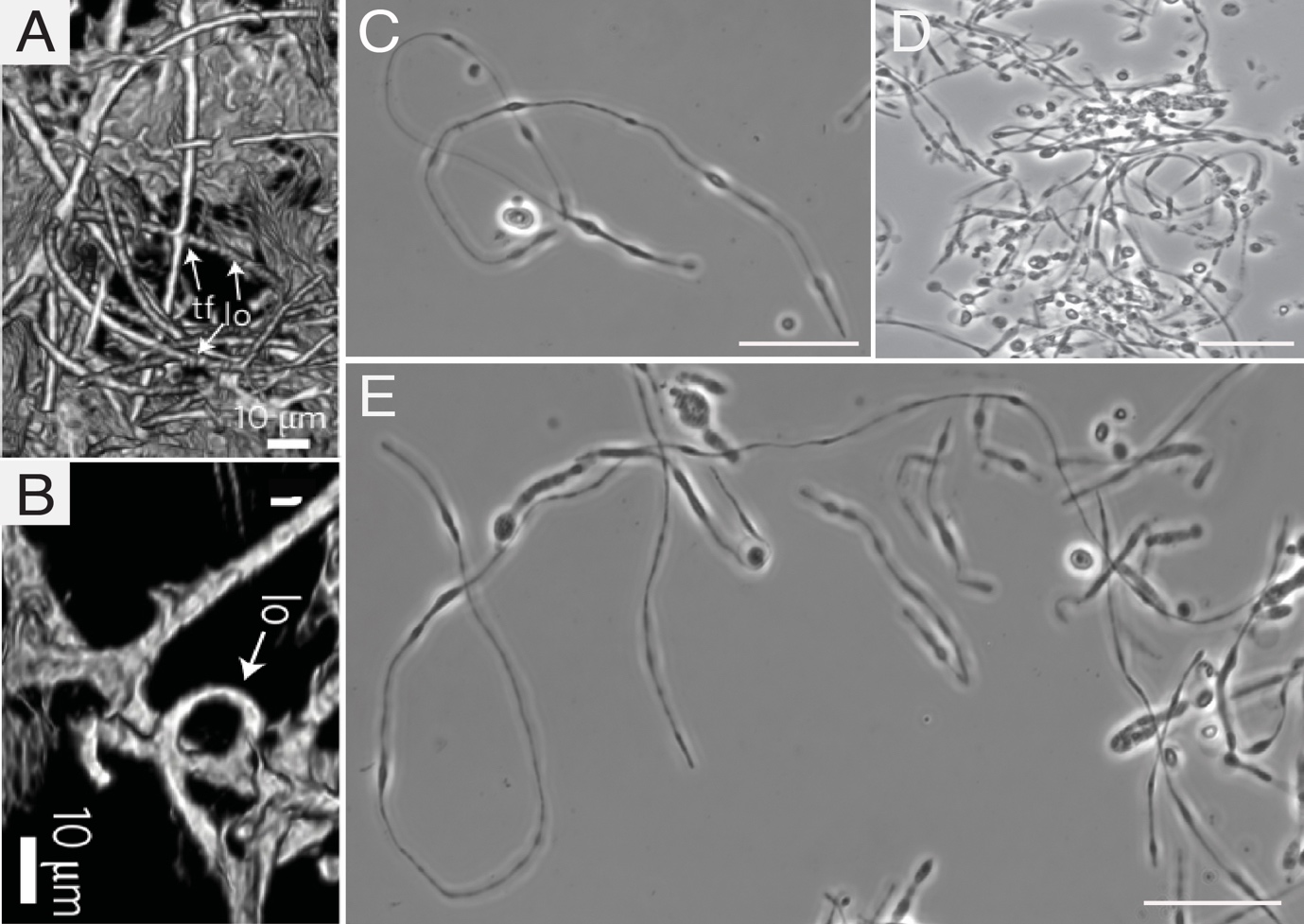

**Fig S3. Morphological comparison of Ongeluk formation microfossils with *RS-P*.**

Images A & B are microfossils reported from Ongeluk formation. Image C-E is a Phase-contrast microscope image of *RS-P* (Scale bar: 10 µm). Arrows (lo) in images A & B point to loops formed by the filamentous microfossils. Similar looping of cells can be seen in *RS-P*.

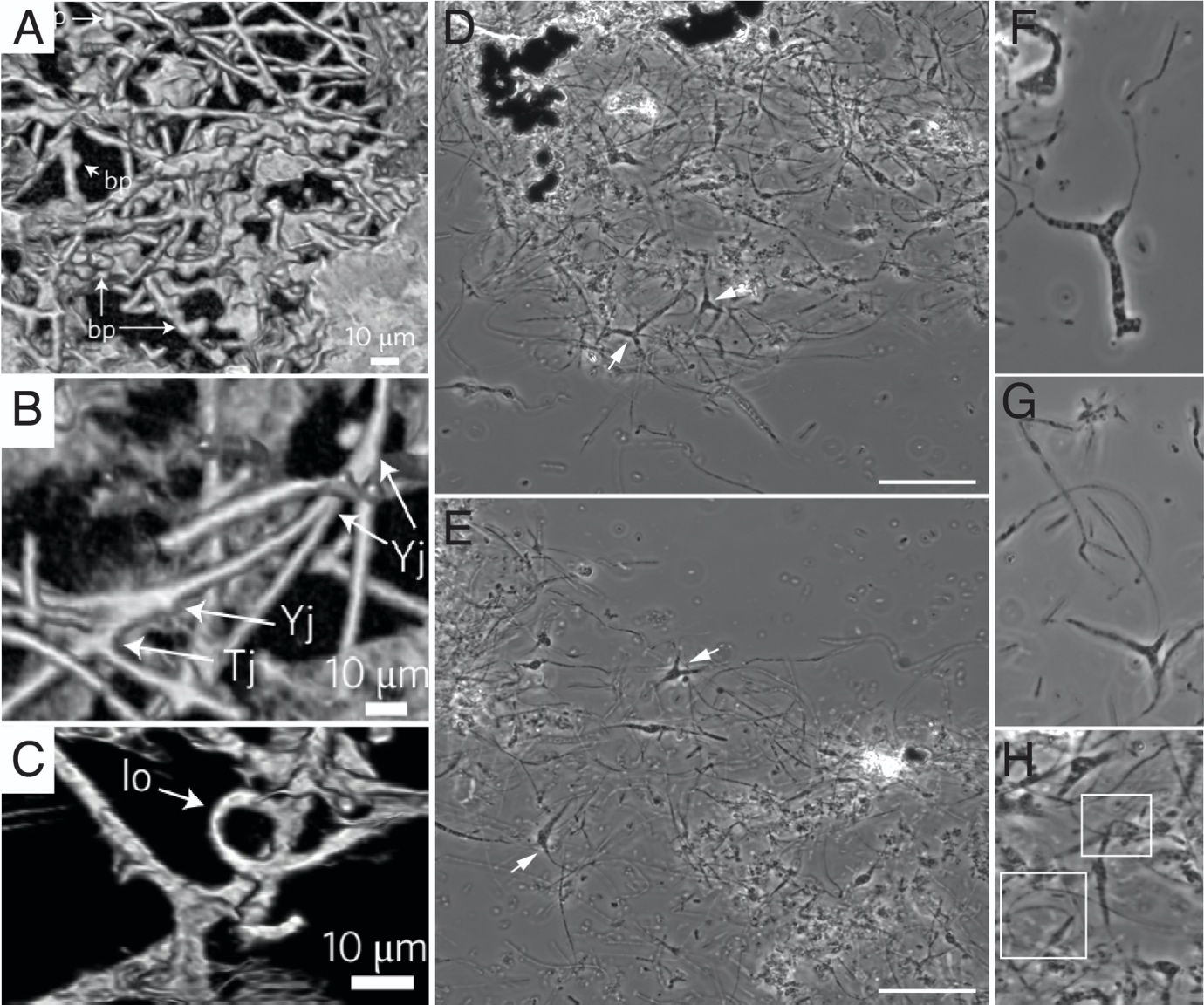

**Fig S4. Morphological comparison of Ongeluk formation microfossils with *RS-P*.**

Images A-C are microfossils reported from Ongeluk formation. Images D-H are phase-contrast microscope images of *RS-P* (Scale bar: 10 µm). Arrows (Yj) in images A-C point to branching within filamentous cells. Arrows in images D-H point to a similar branching pattern in *RS-P*. T-type branching of *RS-P* filaments was shown in Fig S5.

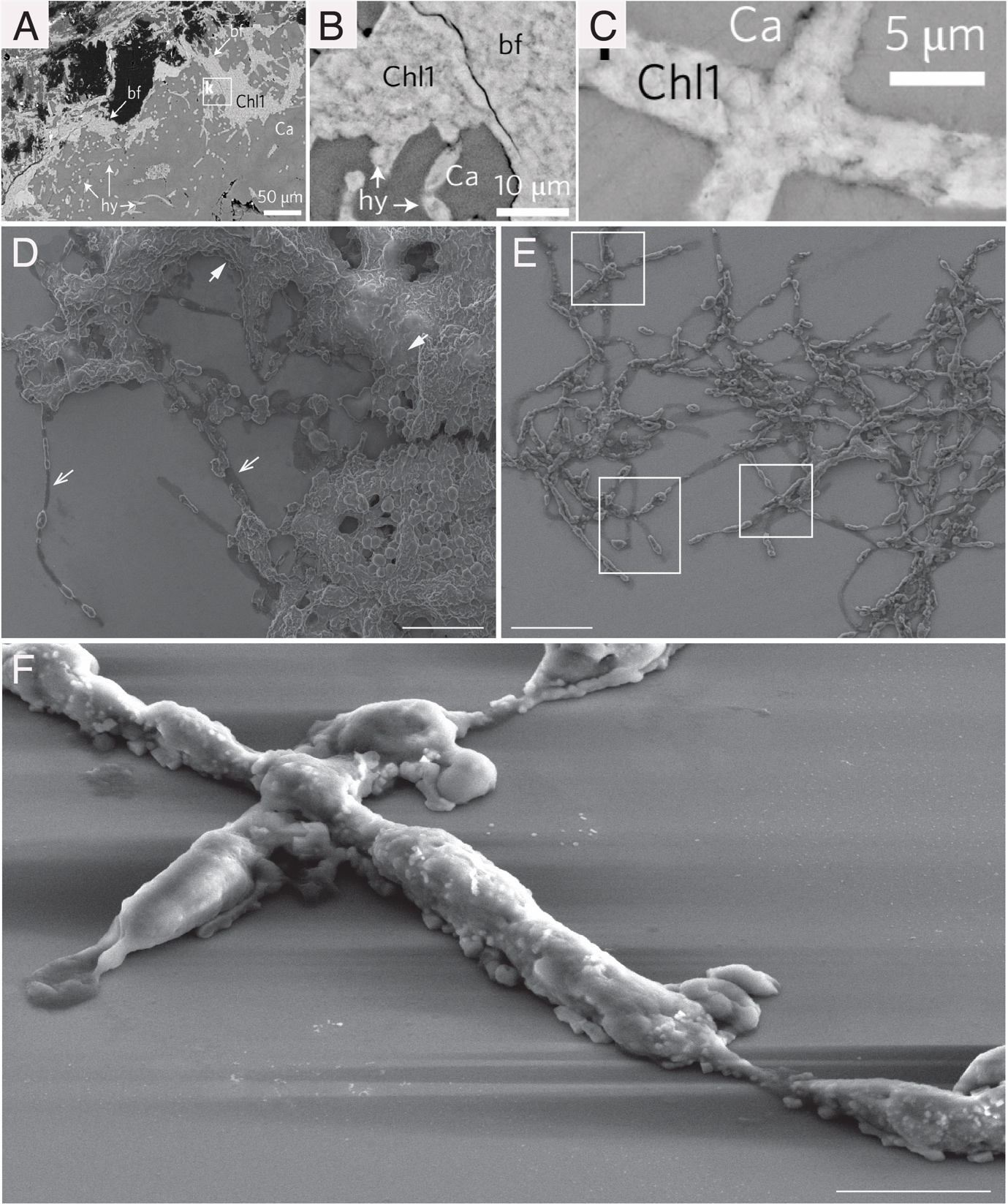

**Fig S5. Morphological comparison of Ongeluk formation microfossils with *RS-P*.**

Images A-C are microfossils reported from Ongeluk formation. Images D-F are Scanning electron microscope images of *RS-P* (Scale bar: 2 µm). Arrows (hy) in images A-C points to hyphae-like structures (bf – biofilm & Chl1 – Chlorite 1). Open arrows in image D point to hollow and collapsed regions within *RS-P*. Closed arrows in these images point to mineral-encrusted cells of *RS-P*, similar in their morphology to cells shown in images A&B. Boxed regions within image E show the juxtaposed filamentous cells. Image F also shows juxtaposed cells forming a T-junction, like the Ongeluk formation microfossils shown in image C.

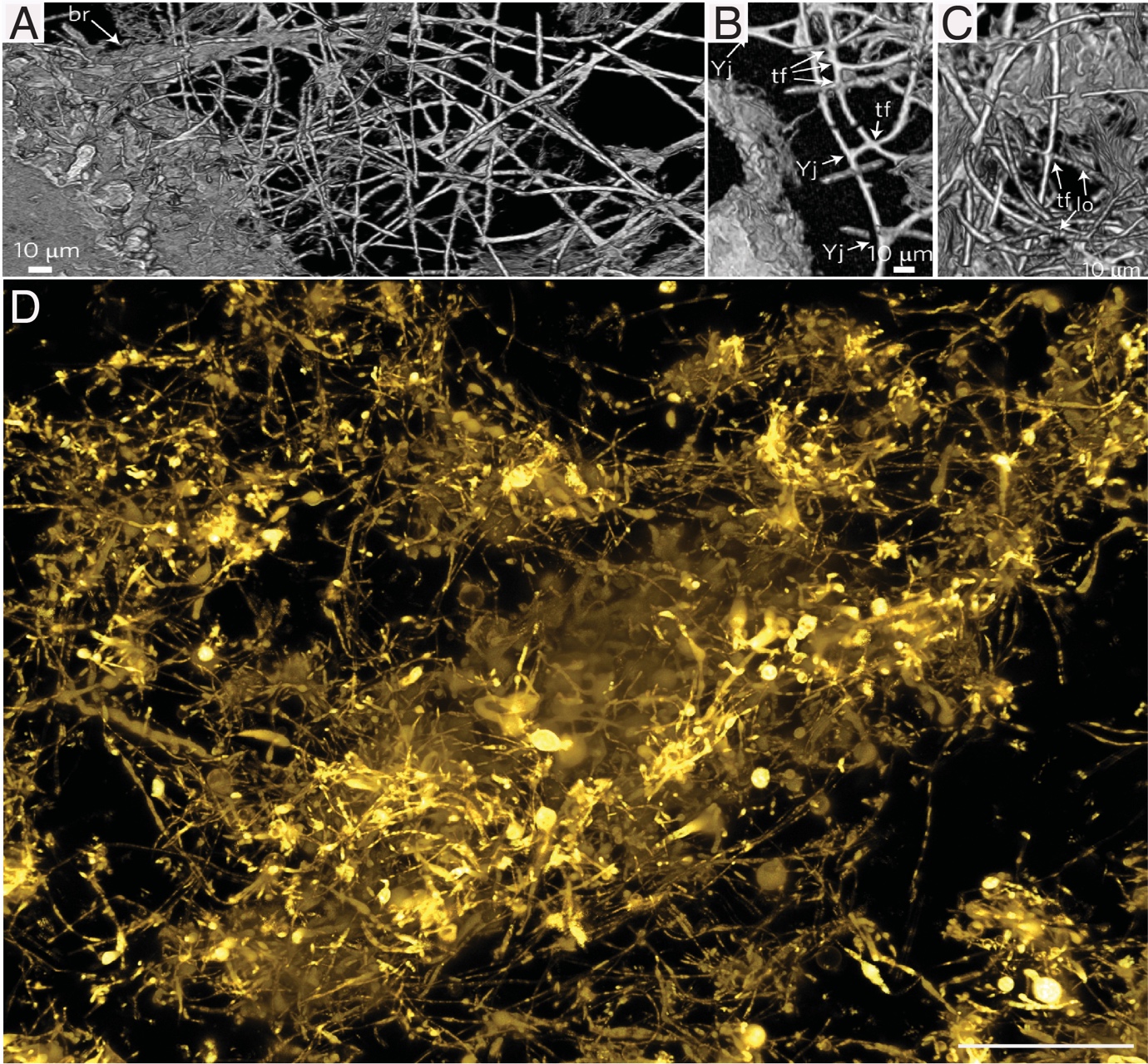

**Fig S6. Morphological comparison of Ongeluk formation microfossils with *RS-P*.**

Images A-C are microfossils reported from Ongeluk formation. Image C is a confocal microscope image of *RS-P* (Scale bar: 20 µm). Cells in these images were stained with universal membrane stain, FM^TM^5-95 (yellow). The clumping of *RS-P* cells can be seen in the image. Subsequent lysis of these cells and the formation of membrane debris, as shown in Fig S7.

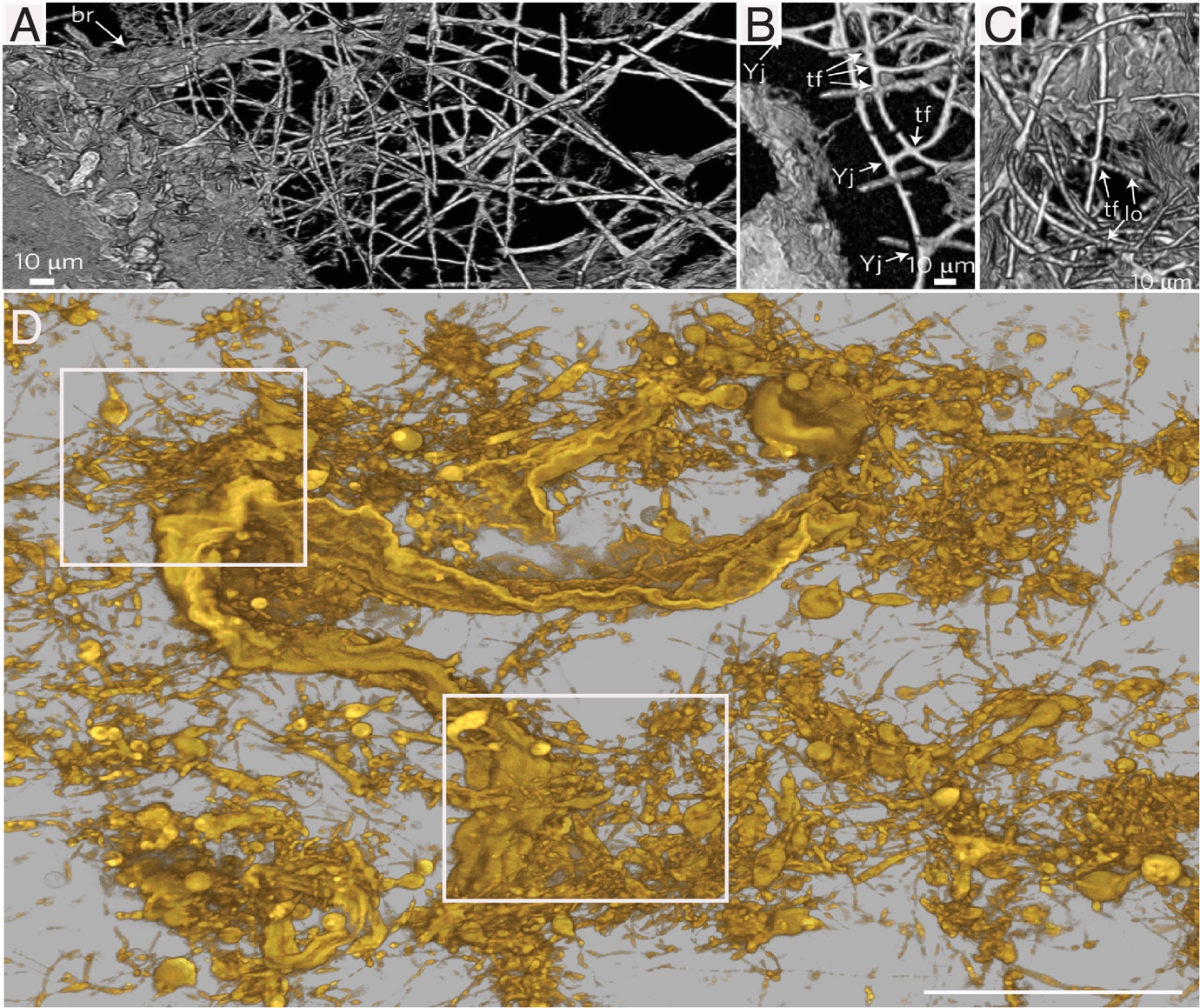

**Fig S7. Morphological comparison of Ongeluk formation microfossils with *RS-P*.**

Images A-C are microfossils reported from Ongeluk formation. Image C is a confocal microscope image of *RS-P* (Scale bar: 20 µm). Cells in these images were stained with universal membrane stain, FM^TM^5-95 (yellow). Cells in these images are closely associated with membrane debris formed from the cell lysis.

**Fig S8. Morphological comparison of Gunflint iron formations microfossils with *RS-P*.**

Images A&B are microfossils reported from Gunflint iron formations. Images C&D are phase-contrast images of morphologically similar *RS-P*’s membrane debris (Scale bar: 10 µm). Sequential morphological transformation of tubular and bulbous *RS-P* cells leading to the formation of these structures was shown in Fig S9 & S10.

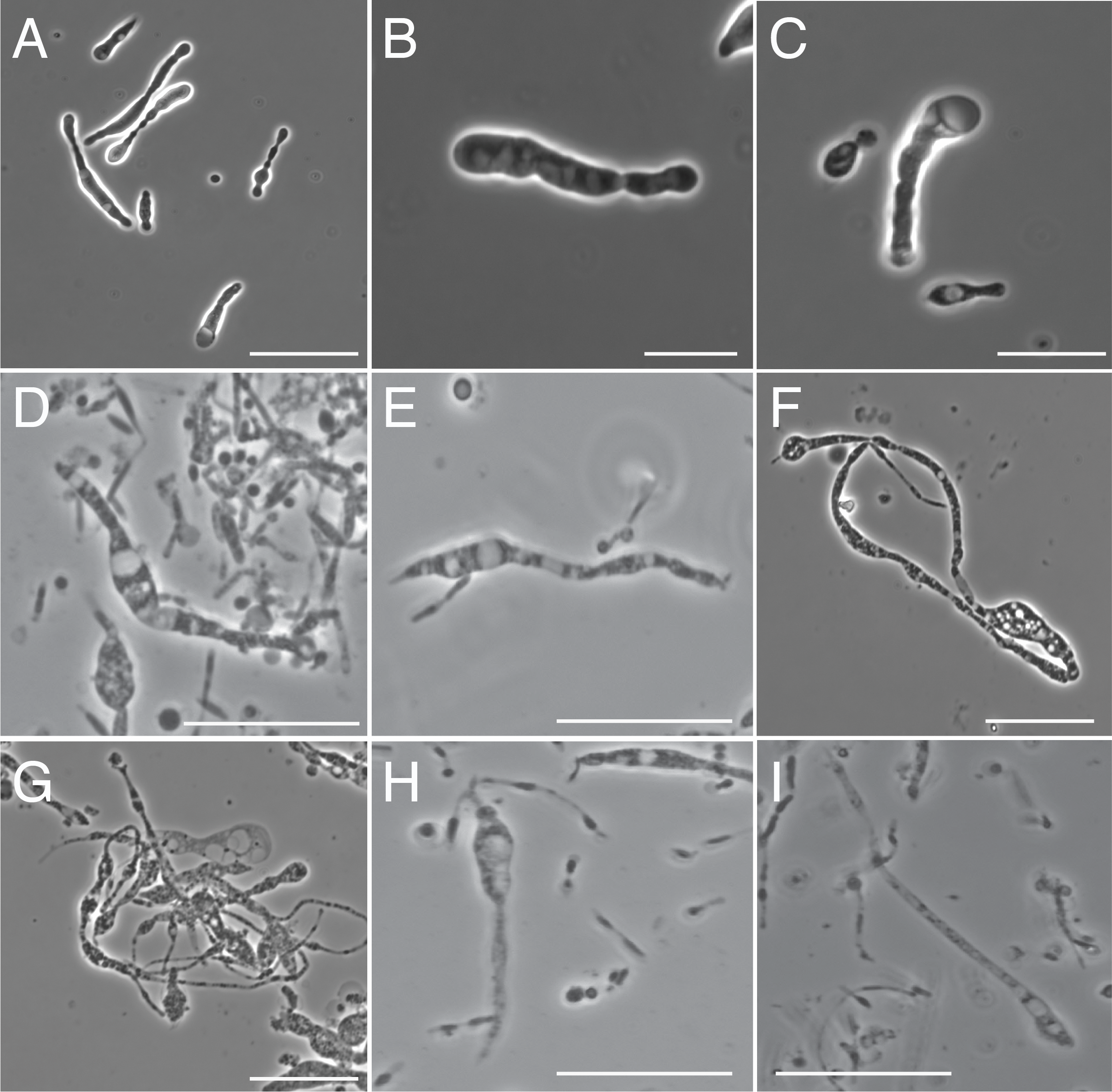

**Fig S9. Sequential growth stages of tubular *RS-P* cells.**

Images A-I are phase-contrast images depicting the sequential morphological transformation of tubular *RS-P* leading to the formation of membrane debris with internal septa, like the microfossils reported from Gunflint iron formations (Scale bar: 10 µm).

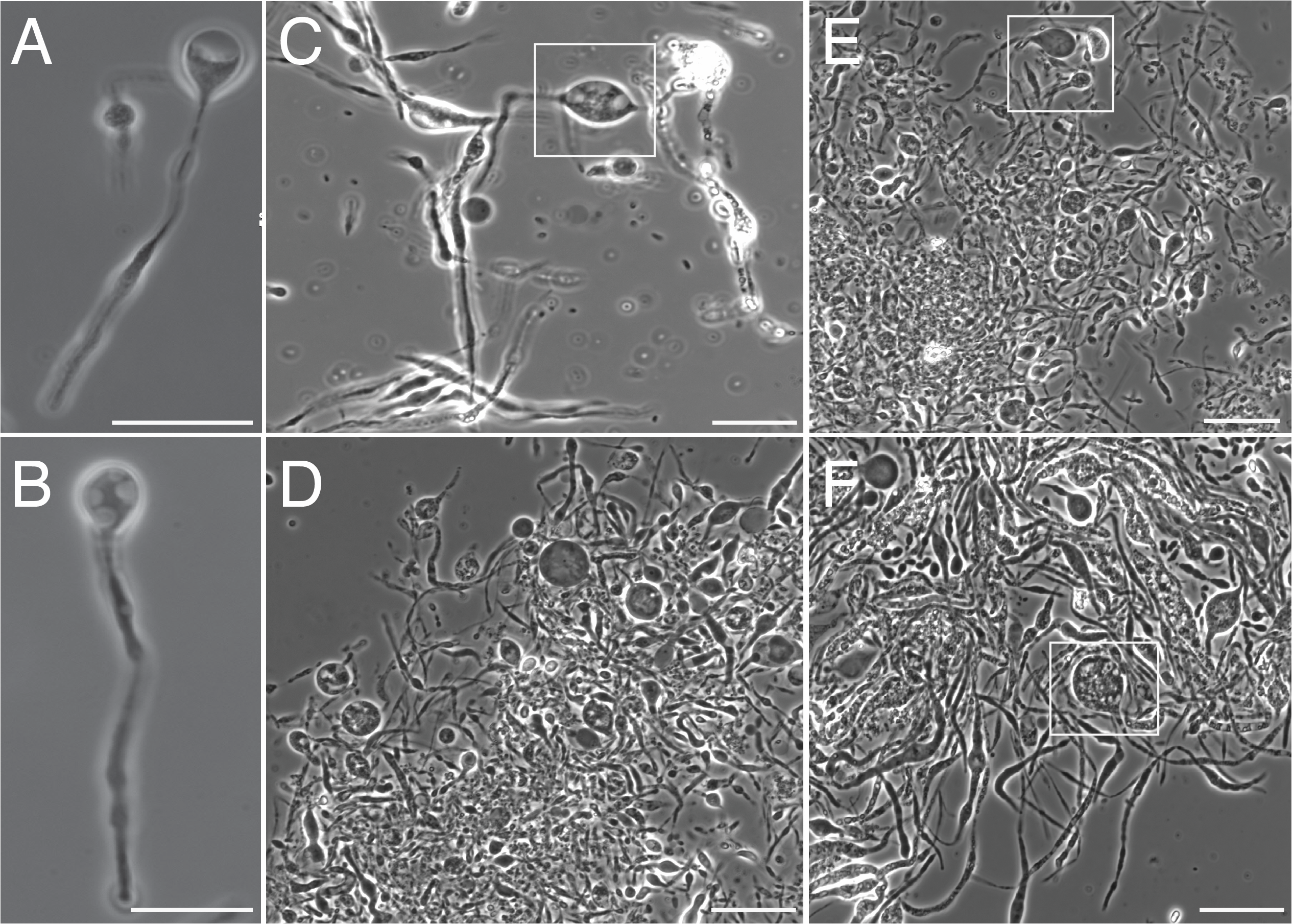

**Fig S10. Sequential growth stages of bulbous *RS-P* cells.**

Images A-F are phase-contrast images depicting the sequential morphological transformation of tubular *RS-P* leading to the formation of membrane debris with internal septa, like the microfossils reported from Gunflint iron formations (Scale bar: 10 µm). Boxed regions in images C, D & F show bulbous cells with internal septa.

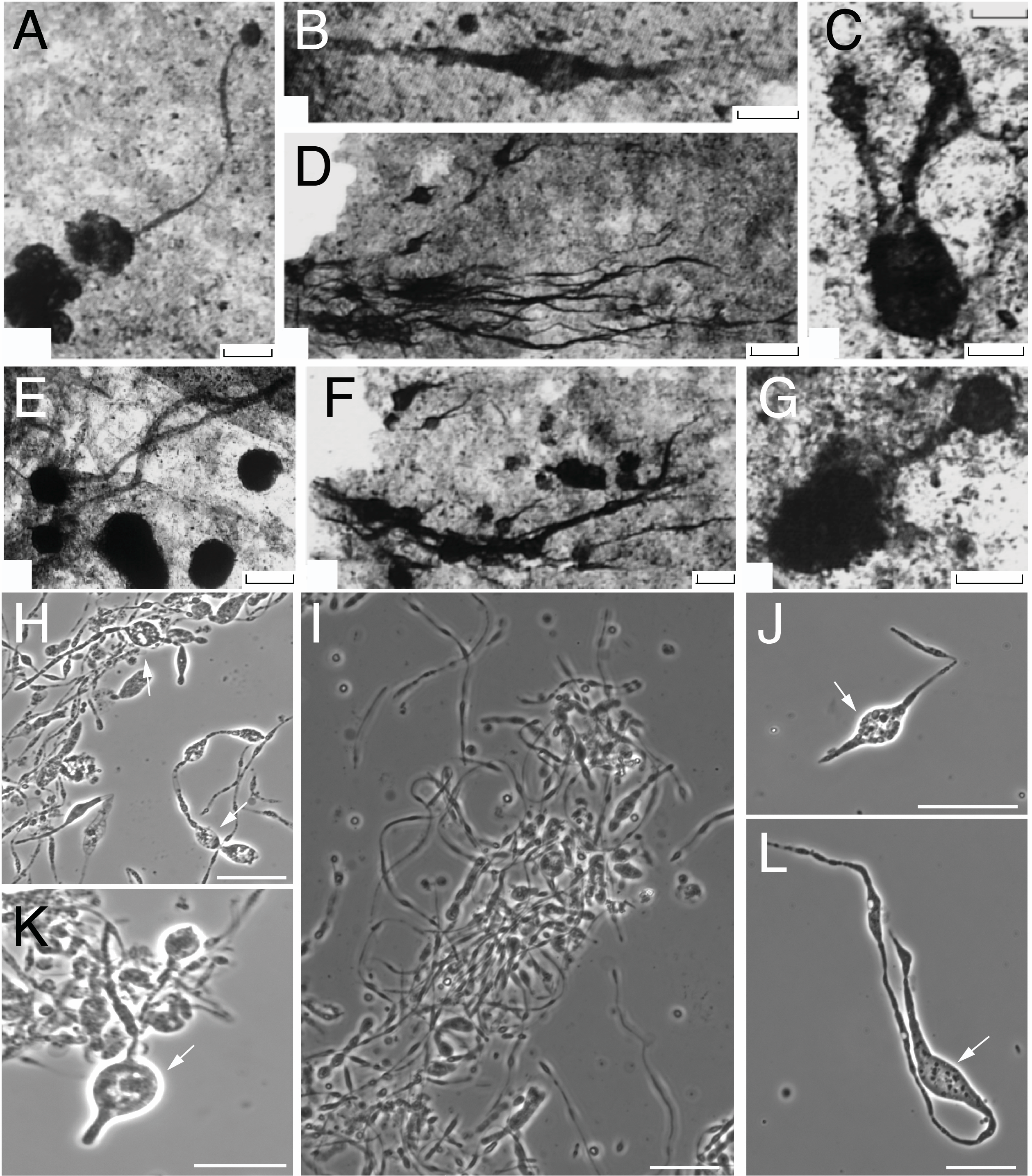

**Fig S11. Morphological comparison of Lakhanda formation microfossils with *RS-P*.**

Images A-G are microfossils reported from the Lakhanda formation. These microfossils resemble fungal zygospores, bulbous structures within filamentous cells. Scale bars: A, B & E: 50 µm, C & G: 20 µm, D & F: 100 µm. Images H-L are phase-contrast microscope images of *RS-P* exhibiting similar morphologies. Arrows in these images point to bulbous structures within filamentous cells. Some bulbous structures even contain internal vesicles resembling spores within zygospore conidia (Scale bar: 10 µm). Sequential stages involved in their formation are shown in Fig S13.

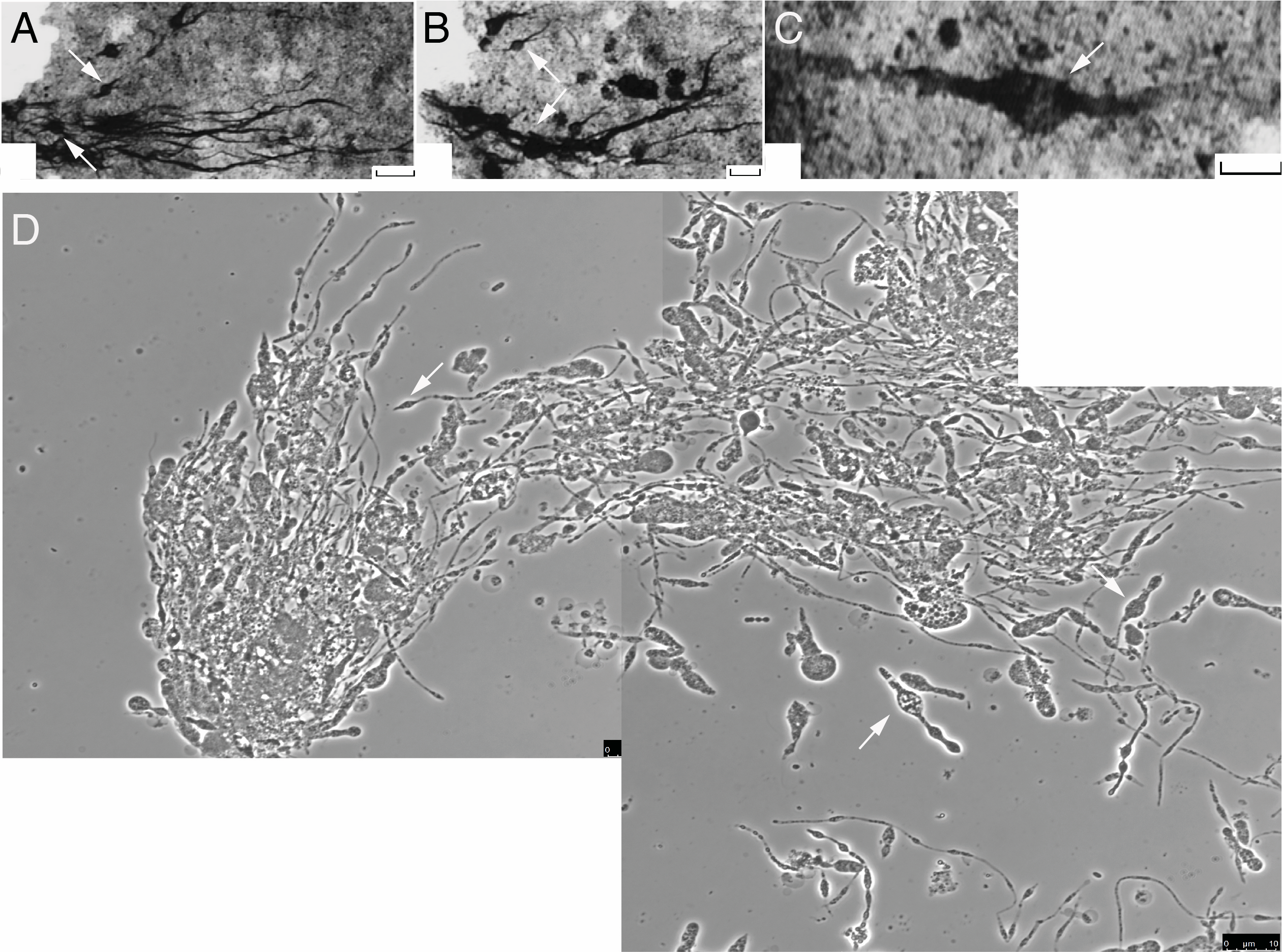

**Fig S12. Morphological comparison of the Lakhanda formation microfossils with *RS-P*.**

Images A-C are microfossils reported from the Lakhanda formation. These microfossils resemble fungal zygospores, bulbous structures within filamentous cells. Scale bars: A&B: 100 µm, C: 50 µm. Image D is phase-contrast microscope images of *RS-P* exhibiting similar morphologies. Arrows in these images point to bulbous structures within filamentous cells. Some bulbous structures even contain internal vesicles resembling spores within zygospore conidia (Scale bar: 10 µm). Sequential stages involved in their formation are shown in Fig S13.

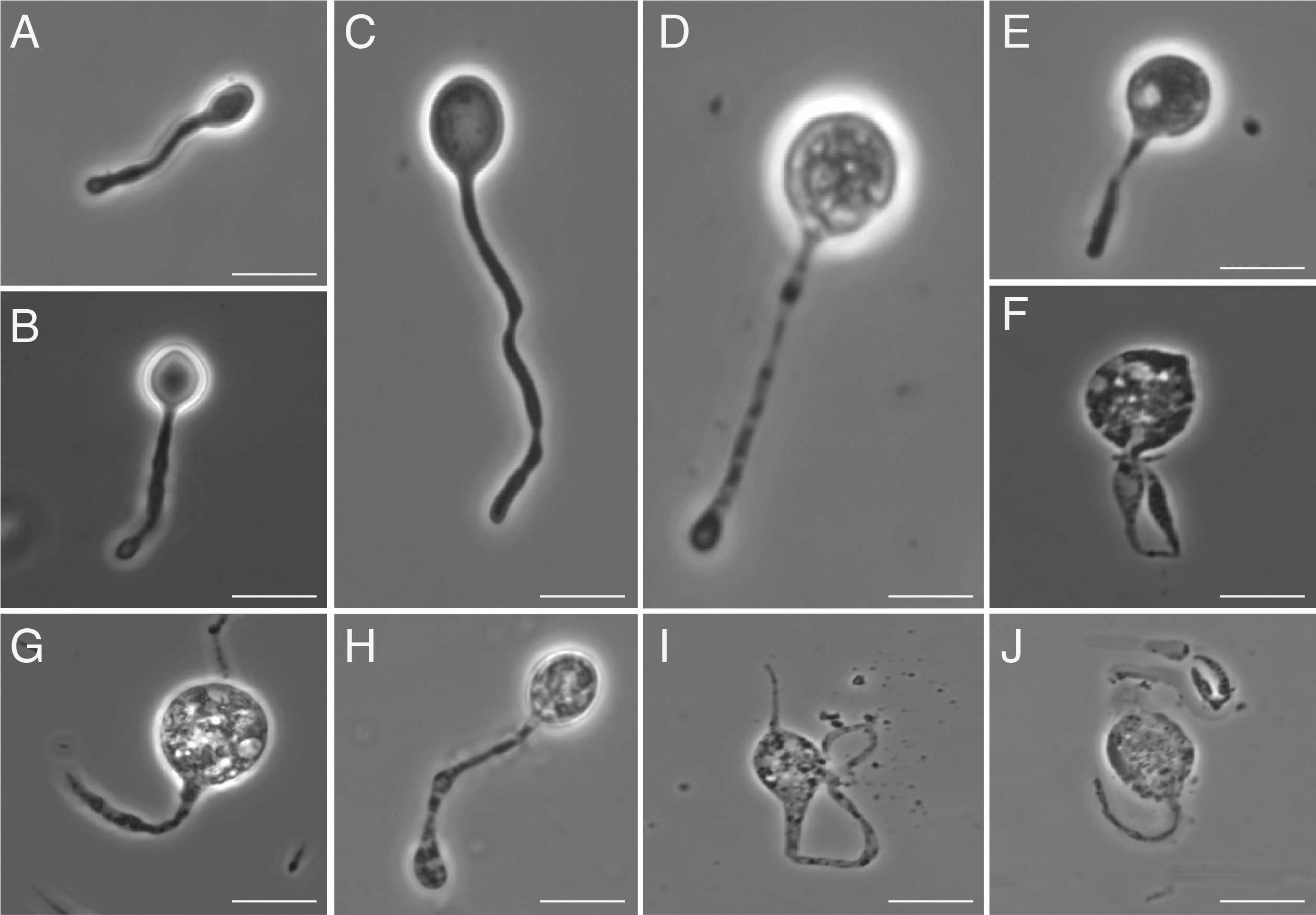

**Fig S13. Sequential growth stages of bulbous conidia-like structures of *RS-P*.**

Images A-J are phase-contrast images of *RS-P*’s bulbous conidia-like structures (scale Bars: 10 µm). Images A-C show early growth stages with uniformly distributed cytoplasm. Images D-G show cells forming internal membrane vesicles, giving the cells a reticulate appearance. Images H-I are the last growth stage cells undergoing lysis.

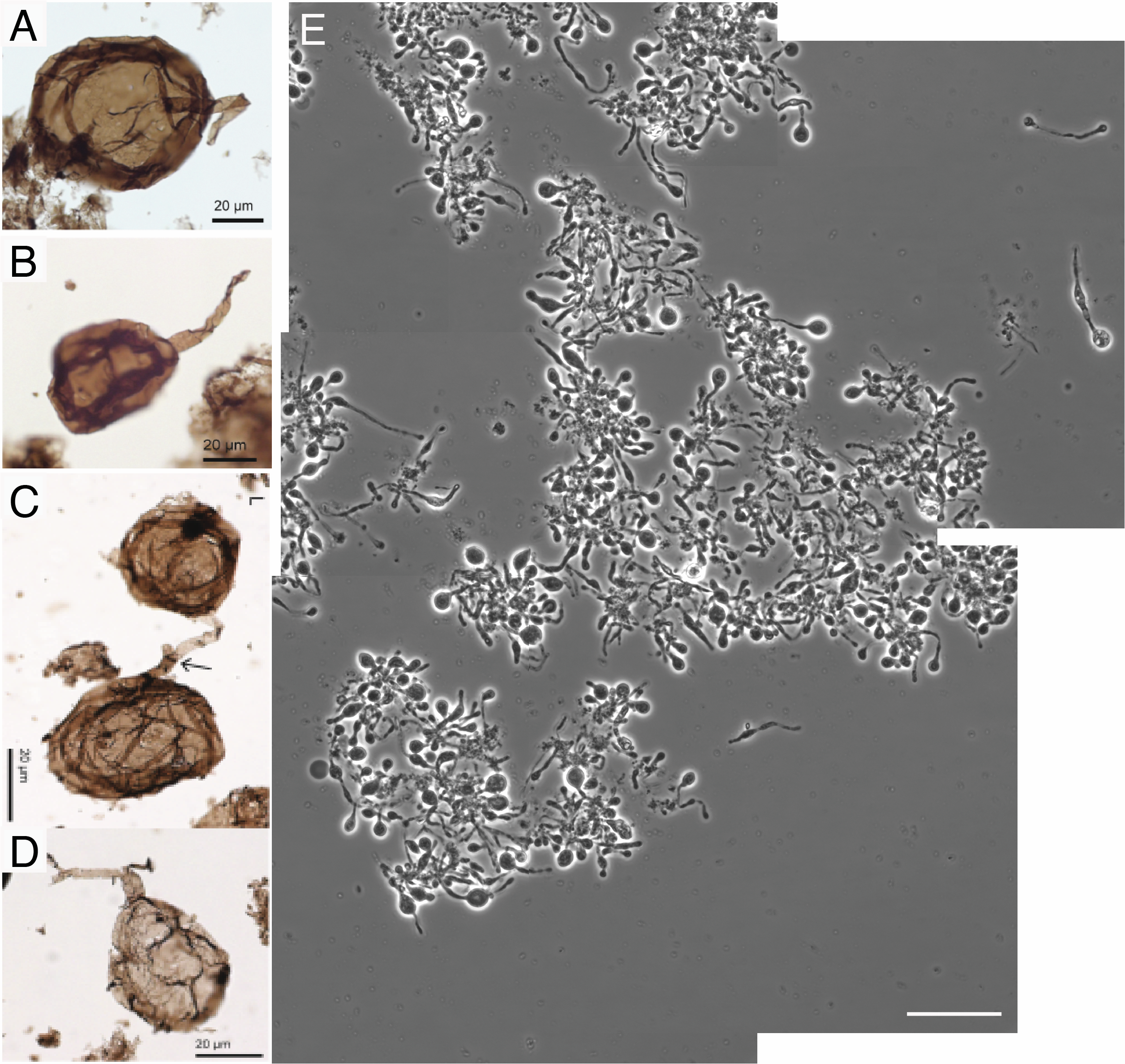

**Fig S14. Morphological comparison of Grassy Bay formation microfossils with *RS-P*.**

Images A-D are microfossils reported from the Grassy Bay formation. Image E is a composite phase-contrast image of early growth stage *RS-P* (Scale bar: 10 µm).

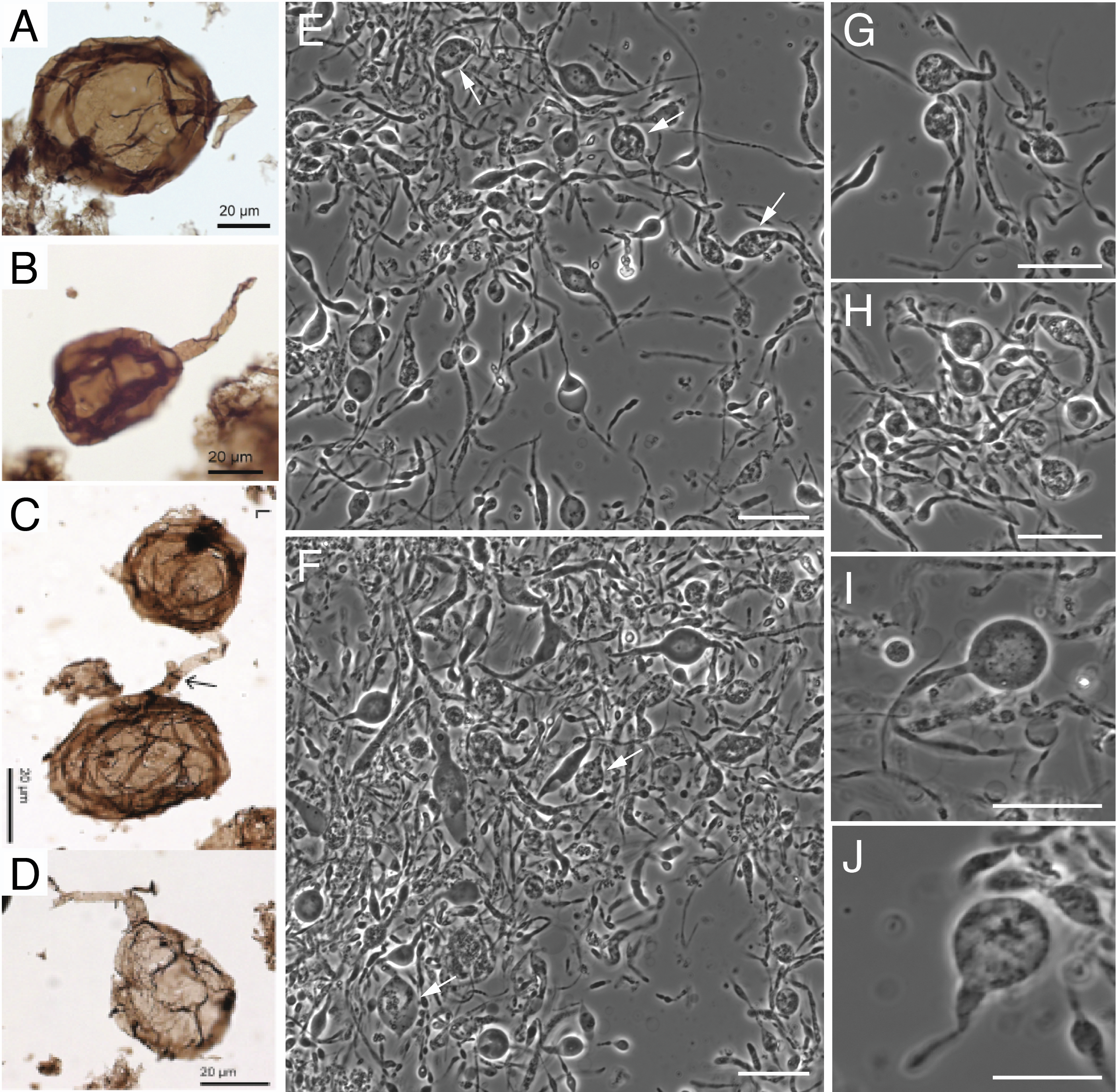

**Fig S15. Morphological comparison of Grassy Bay formation microfossils with *RS-P*.**

Images A-D are microfossils reported from the Grassy Bay formation. Images E-J are phase-contrast images of late-growth stage *RS-P* cells (Scale bar: 10 µm). Arrows in images E&F points to fungal conidia-like structures formed by *RS-P*. Reticulate interiors and membrane vesicles within these cells resemble spores within the fungal conidia.

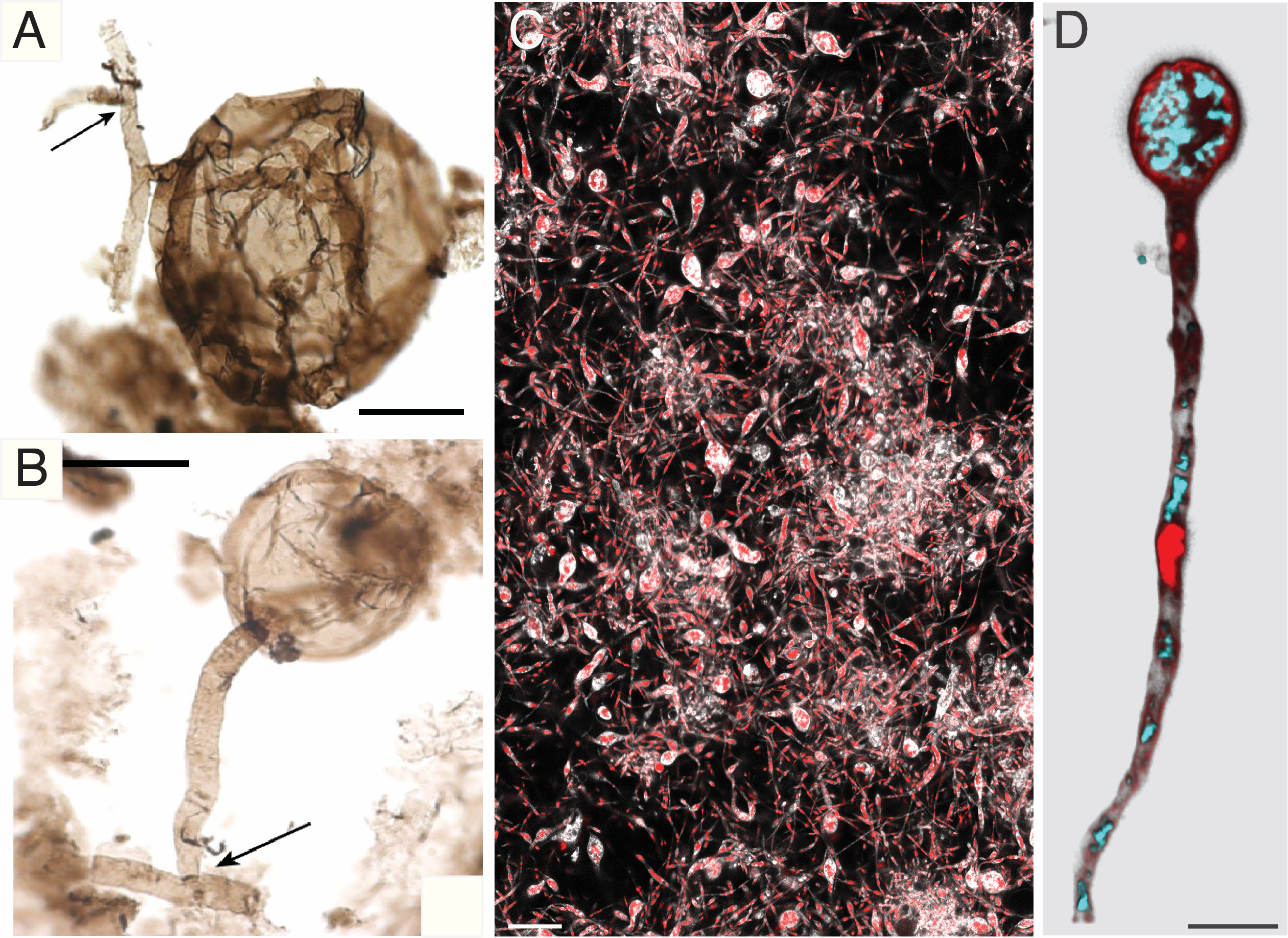

**Fig S16. Morphological comparison of Grassy Bay formation microfossils with *RS-P*.**

Images A&B are microfossils reported from the Grassy Bay formation. Image C&D is a confocal microscope image of *RS-P* (Scale bar: 10 µm). Cells in this image-C are stained with universal membrane stain, FM^TM^5-95 (white), and DNA stain, PicoGreen^TM^ (red). Cells in this image-D are stained with universal membrane stain, FM^TM^5-95 (white), and, L_d_-membrane stain, FAST-Dil^TM^ (Cyan). Reticulate interiors and membrane vesicles within these cells resemble spores within the fungal conidia.
